# Towards the Target: Tilorone, Quinacrine and Pyronaridine Bind to Ebola Virus Glycoprotein

**DOI:** 10.1101/2020.05.26.118182

**Authors:** Thomas R. Lane, Sean Ekins

## Abstract

Pyronaridine, tilorone and quinacrine were recently identified by a machine learning model and demonstrated *in vitro* and *in vivo* activity against Ebola virus (EBOV) and represent viable candidates for drug repurposing. These drugs were docked into the crystal structure of the ebola glycoprotein and then experimentally validated *in vitro* to generate Kd values for tilorone (0.73 μM) pyronaridine (7.34 μM), and quinacrine (7.55 μM). These are more potent than the previously reported toremifene (16 μM).

---

The ongoing Ebola virus (EBOV) outbreak in the Democratic Republic of the Congo, has killed over 2200 people at the time of writing according to the World Health Organization (1). While there is an approved vaccine for prevention of EBOV disease (2) we have no approved antivirals to treat patients although there are several treatments that have reached the clinic (3-6) with the most promising results for biologics only. One target of particular interest for antibody therapies (7) is the glycoprotein which is composed of GP1 and GP2 subunits and is involved in attachment to the cell and entry (8). Earlier high throughput screens had identified benzodiazepine analogs as GP1 inhibitors which act early in viral entry (8, 9). Several structurally diverse FDA approved drugs have also been identified including toremifene, benztropine, bepridil, paroxetine, sertraline and ibuprofen (10) which all bind in the same site at the tunnel entrance to the glycoprotein, destabilizing it and resulting in release of GP2, thus preventing fusion between virus and endosome (9). Several other research groups have identified small molecules that inhibit glycoprotein, suggesting a wide array of molecules may bind (11-15). Others have used computational approaches to perform virtual screens to identify inhibitors of GP2 (16). Recent efforts to identify small molecules drugs for EBOV have included using computational methods in the form of a Bayesian machine learning (ML) approach trained with EBOV inhibitors (17, 18). This ML model enabled a virtual screen and selection of three compounds, tilorone, quinacrine and pyronaridine tetraphosphate (19). All these drugs inhibited EBOV *in vitro* and *in vivo* in the mouse-adapted EBOV (ma-EBOV) efficacy model (20-23). Pyronaridine tetraphosphate also demonstrated significant activity in the guinea pig adapted model of EBOV infection (24) and is currently used as an antimalarial in combination with artesunate (Pyramax®). Most recently pyronaridine was shown to be a potent lysosomotropic agent while artesunate which is also active against EBOV *in vitro* was not (25). Combining these two drugs had an additive effect on inhibiting EBOV replication *in vitro* and reduced cytotoxicity (25). Pursuit of the potential target of these computationally identified drugs pointed to docking compounds in the glycoprotein structure and experimental validation.

## Pyronaridine, tilorone and quinacrine are predicted to bind to EBOV glycoprotein

Our three EBOV inhibitors: tilorone, pyronaridine and quinacrine were docked into the EBOV glycoprotein structure (PDB 5JQ7) at the same site as previously described for toremifene (10) using Discovery Studio LibDock. Binding energies were calculated following a ligand minimization (rigid protein). As a control, toremifene had a calculated minimized binding energy of −114.772 kcal/mol. Pyronaridine had the best score −277.63 kcal/mol, followed by tilorone (−171.73 kcal/mol) and quinacrine (−164.07 kcal/mol) (Fig 1).

**Figure 1.**
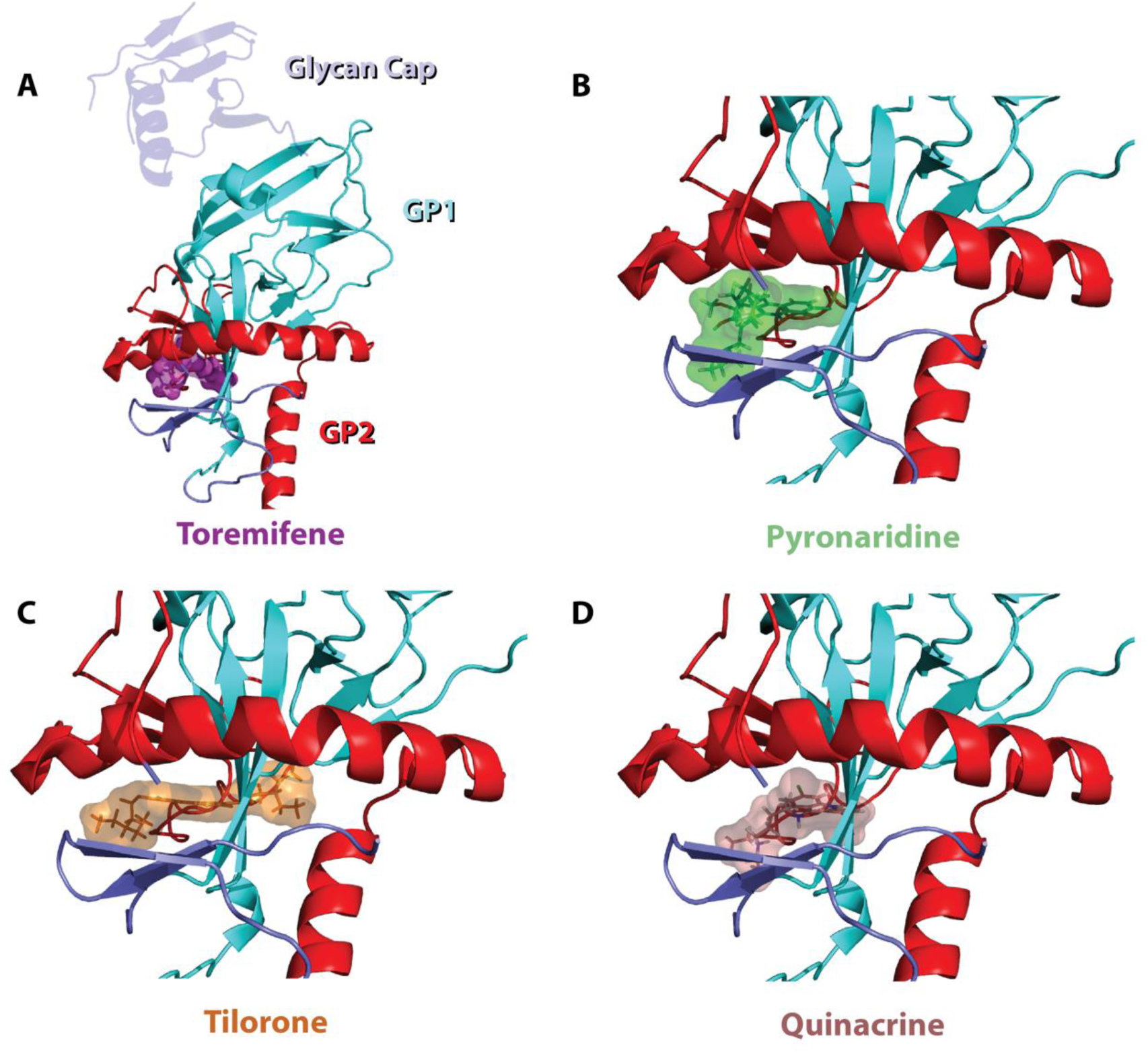
Docking of ligands in EBOV glycoprotein crystal structure showing lowest energy poses from the rigid docking (libdock). Cartoon representations of EBOV glycoprotein (GP1(Cyan)-GP2(Red)) with a glycan cap (purple). A) Crystal structure of EBOV GP in complex with toremifene (PDB ID: 5JQ7; 2.69 Å) As a control, toremifene had a calculated minimized binding energy of −114.772 kcal/mol. Top scoring docked poses of pyronaridine (B, −277.63 kcal/mol), tilorone (C, −171.73 kcal/mol) and quinacrine (D, −164.07 kcal/mol) in EBOV GP using libdock. Binding energies were calculated following a ligand minimization (rigid protein).

## Pyronaridine, tilorone and quinacrine bind to EBOV glycoprotein *in vitro*

Pyronaridine tetraphosphate [4-[(7-Chloro-2-methoxybenzo[b][1,5]naphthyridin-10-yl)amino]-2,6-bis(1-pyrrolidinylmethyl)phenol phosphate (1:4)] (19) was purchased from BOC Sciences (Shirley NY). Tilorone and quinacrine were purchased from Cayman Chemicals (Ann Arbor, Michigan). Toremifene and clomiphene were purchased from MedChemExpress (Monmouth Junction, NJ) and favipirivir from TRC Canada (North York, ON Canada). Zaire ebolavirus disulfide-linked glycoprotein heterodimer (GP1-GP2) was purchased from Novus biologicals (Centennial, CO). According to the manufacturer, EBOV GP protein is purified from CHO-derived viral expression with previous internal verification of significant glycosylation. 200 µg of lyophilized protein was resuspended in RED-NHS 2nd Generation labeling buffer (NanoTemper; Cambridge, MA). This was followed by the labeling of the primary amines using the RED-NHS dye according to the manufacturers protocol. Labeled protein was buffer exchanged into 10 mM MES, pH 5.0, 150 mM NaCl, 170 mM sodium malonate at pH 5.2 (MST buffer), which is similar to a buffer previously shown to be appropriate for EBOV GP (10), and then diluted to a final concentration of 1 µM. For each experimental compound 16 independent stocks were made in DMSO using 2-fold serial dilution (10 mM initial concentration). The MST buffer used for a final dilution prior to MST was supplemented with 0.05% Tween 20 and 10 mM BME. The protein was diluted to 2.5 nM in the supplemented MST buffer and 19.5 µL of this was combined with 0.5 µL of the compound stock and then mixed thoroughly. This resulted in 2-fold serial dilution testing series with the highest and lowest concentration of 250 µM and 7.629 nM, respectively, with a consistent final DMSO concentration of 2.5%. These reactions were incubated for 20-30 min prior to transferring to standard Monolith NT.115 capillaries. Experiments were run at 20% excitation and high MST power at 23.0°C on a Monolith NT.115Pico (NanoTemper). Favipiravir and toremifene were also run as the negative and positive controls, respectively. Each experimental compound was run in quadruplicate.

## Data acquisition and analysis

The data were acquired with MO.Control 1.6.1 (NanoTemper Technologies). Recorded data were analyzed with MO.Affinity Analysis 2.3 (NanoTemper Technologies). The dissociation constant Kd quantifies the equilibrium of the reaction of the labelled molecule A (concentration c_A_) with its target T (concentration c_T_) to form the complex AT (concentration c_AT_): and is defined by the law of mass action as: 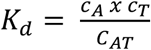, where all concentrations are “free” concentrations. During the titration experiments the concentration of the labelled molecule A is kept constant and the concentration of added target T is increased. These concentrations are known and can be used to calculate the dissociation constant. The free concentration of the labelled molecule A is the added concentration minus the concentration of formed complex AT. The Kd is calculated as 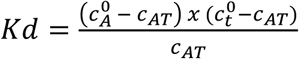. The fraction of bound molecules x can be derived from F_norm_, where F_norm_(A) is the normalized fluorescence of only unbound labelled molecules A and F_norm_(AT) is the normalized fluorescence of complexes AT of labeled as shown by the equation: 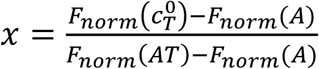. The MST traces that showed aggregation or outliers were removed from the datasets prior to Kd determination.

The Kd values for pyronaridine (7.34 μM), tilorone (0.73 μM) and quinacrine (7.55 μM) were lower than for the positive control toremifene (24.83 μM) which is very similar to the literature value of 16 µM run under similar conditions (10). Additionally, we identified clomiphene (30.74 μM), whereas favipiravir and artesunate did not bind to the EBOV glycoprotein (Figure 2). Favipiravir is an RNA polymerase inhibitor and would not be expected to bind to the glycoprotein. We have previously demonstrated that artesunate does not inhibit lysotracker (25) and is likely inhibiting ebola via a different mechanism to pyronaridine with which it is used in combination as the drug Pyramax™ for malaria. The lack of inhibition of glycoprotein by artesunate provides further confirmation of this. Prior work on pyronaridine has demonstrated that it has promising *in vitro* activity against EBOV as well as excellent *in vitro* absorption, distribution, metabolism and excretion (ADME) properties and a very long half-life across mice and humans (21). It also has statistically significant *in vivo* efficacy in mice and guinea pigs infected with EBOV (24). While the mechanism to date was unclear, it has been shown to inhibit lysostracker suggesting a lysosomotropic effect (25). When this is coupled with the potential to bind to glycoprotein this dual mechanism will likely prevent EBOV entry. Similarly, quinacrine, is a small-molecule, orally bioavailable drug which has also been used clinically as an antimalarial. This also has very favorable ADME properties (apart from potent CYP2D6 inhibition) and demonstrated *in vivo* activity in mice infected with EBOV (20). Tilorone is structurally different and is used in eastern Europe as an antiviral. It has excellent *in vitro* ADME properties and was recently demonstrated to have potent *in vitro* inhibition of EBOV as well as efficacy in mouse infected with EBOV (22). Interestingly this molecule did not demonstrate efficacy in the guinea pig EBOV model likely due to the significant species differences in metabolism when compared to mouse (24). Tilorone has also demonstrated *in vitro* inhibition of MERS (26) and is under evaluation as a potential treatment for other coronaviruses such as SARS-CoV-2 (23) and has shown a low μM IC_50_ (27). This current work now provides further detail as to how these molecules target EBOV glycoprotein as well as being lysosomotropic, enabling their blocking of viral entry. This information may aid in structure guided modification of these compounds and future X-ray crystallography. Such mechanistic insights may aid drug discovery for other viruses (e.g. SARS-CoV-2). To date tilorone would appear the highest affinity compound for EBOV glycoprotein compared to the previously reported toremifene (10, 15).

**Figure 2.**
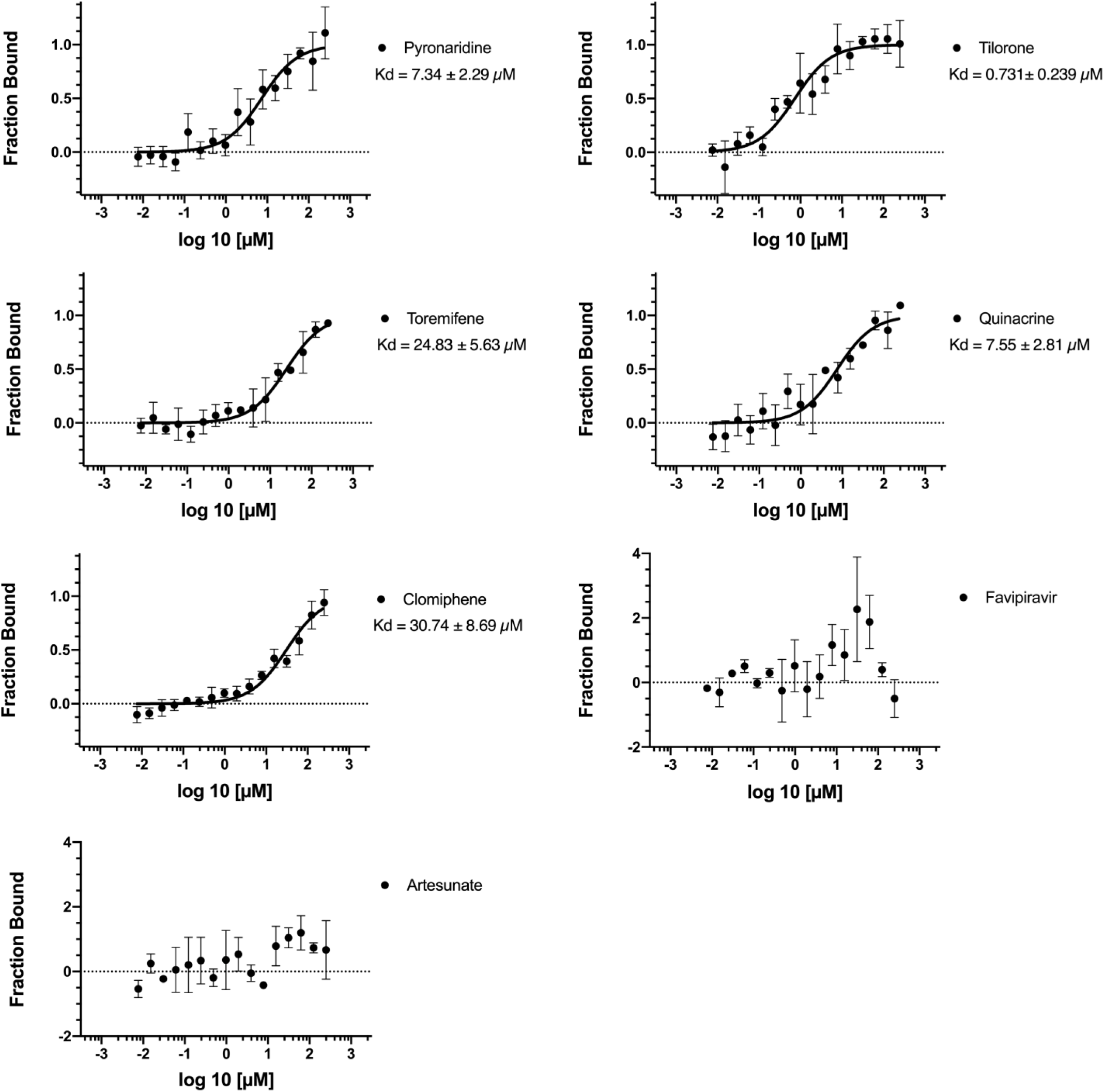
Ebola glycoprotein Kd values generated using microscale thermophoresis for test compounds.

In conclusion, the accumulated *in vitro* and *in vivo* data gathered for tilorone, quinacrine and pyronaridine points to them sharing a common target or mechanism for the inhibition of EBOV (20-22, 24, 25, 28). It would also appear they all block entry, are lysosomotropic and now are identified to bind to the glycoprotein. Shedding further light on how these molecules work *in vitro* may provide further justification for clinically repurposing these compounds.

## ACKNOWLEDGMENTS

We would like to sincerely thank Dr Peter Madrid, Dr. Jason Comer, Dr. Julie Dyall, Dr. Manu Anantpadma, and Dr. Robert Davey for many Ebola related discussions. Dr. Ana Puhl Is gratefully acknowledged for discussions on MST.

